# Integrative analysis of plant responses to a combination of water deficit, heat stress and eCO_2_ reveals a role for OST1 and SLAH3 in regulating stomatal responses

**DOI:** 10.1101/2025.05.07.652739

**Authors:** María Ángeles Peláez-Vico, Ranjita Sinha, Abdul Ghani, María F López-Climent, Trupti Joshi, Felix B. Fritschi, Sara I. Zandalinas, Ron Mittler

## Abstract

The frequency, intensity and duration of global change factors and/or environmental stressors, such as droughts, heat waves, floods, and pollution, are increasing due to anthropogenic activities, subjecting plants to compound conditions of ‘Global Change Factor combination’ (GCFc), or ‘stress combination’. These can have contrasting or additive effects on plant physiological performance and impact different ecosystems and agro-ecosystems worldwide. Using an integrative physiological, genetic, hormonal, and transcriptomic analysis, we studied the response of *Arabidopsis thaliana* (L.) Heynh to a combination of water deficit (WD), heat stress (HS) and/or elevated levels of CO2 (eCO2). Our findings reveal high specificity in plant responses to GCFc. We further reveal that stomatal aperture regulation under conditions of GCFc is controlled by a blend of unique and shared regulators, that together determine the specific aperture size under each different set of GCFc conditions. Under a combination of WD, HS, and eCO2 (WD+HS+eCO2), for example, stomatal closure required the function of nitric oxide, OPEN STOMATA 1 and the S-type anion channel SLAH3, but not the S-type anion channel SLAC1, or many other regulators typically found to control stomatal aperture under less complex conditions, including HS+WD. Our findings highlight the complexity of plant physiological and molecular responses to conditions of stress combination and open the way for future studies of plant responses to stress combination, crucial to our understanding of how GCFc impacts different ecosystems and agro-ecosystems worldwide. In addition, our study provides an initial definition for a ‘stomatal hierarchical stress code’ that could apply to future studies.

## 1. INTRODUCTION

The intensity, number, duration, and overall complexity of environmental stressors, or global change factors (GCFs), are continuously increasing on our planet due to global warming, climate change, and chemical pollution, subjecting plants and microbiomes to different ‘Global Change Factor combinations’ (GCFc; Lesk et al. 2016; Rillig et al. 2019; Alizadeh et al. 2020; Zandalinas et al. 2021; Lesk et al. 2022; Zandalinas and Mittler, 2022; IPCC, 2023; Richardson et al. 2023; Bi et al. 2024; Rhee et al. 2024). As a result, our future crops, trees, and other plants, living within different ecosystems and agro-ecosystems worldwide, would need to grow under different combinations of different abiotic stress conditions that could include heat waves, droughts, floods/waterlogging, ozone, high salinity, extreme soil pH, enhanced pollution, and/or low nutrient stress, under elevated levels of CO2 (eCO2). These could have adverse and/or beneficial effects on different physiological processes (*e.g.,* stomatal aperture regulation and photosynthesis), plant growth, and reproduction. A recent study has shown, for example, that soybean (*Glycine max*) plants subjected to 5 different abiotic stress conditions in an increasing complexity of stress combination conditions (*i.e.,* GCFc, or ‘Multifactorial stress combination’; MFSC) displayed a gradual decrease in stomatal aperture, transpiration, photosynthesis, growth and yield (Peláez-Vico et al. 2024a).

Recent studies suggest that stomata could play a pivotal role in plant responses to stress combination/GCFc (Zandalinas and Mittler, 2022; Sinha et al. 2022, 2023; Jensen et al. 2024; Peláez-Vico et al. 2024a, 2024b; Xu et al. 2025). Stomata are epidermal pore structures, bordered by two guard cells that regulate their aperture (Lawson and Matthews 2020; Clark et al. 2022). They play a cardinal role in plant physiology, metabolism, development, and pathogen/stress responses, and are thought to have evolved during the establishment of plant life on the surface of our planet (Lawson and Matthews 2020; Postiglione and Muday 2020; Hsu et al. 2021; Clark et al. 2022; Waadt et al. 2022; Chen and Torii 2023; Blatt, 2024; Melotto et al. 2024). Stomata control the exchange of CO2, O2, water, and other gaseous compounds and signals between the inner spaces of plants and the atmosphere, directly impacting processes such as photosynthesis, transpiration, respiration, and biotic and abiotic responses (Kollist et al. 2014; Lawson and Matthews 2020; Waadt et al. 2022; Zhang et al. 2022). Stomatal aperture is known to decrease in response to water deficit (WD), elevated eCO2, ozone stress, excess light (EL) stress, waterlogging, wounding (W), and bacterial pathogen infection, and to open in response to heat stress (HS; to allow cooling of plant tissues via transpiration), low CO2 levels, and transition from dark to light (Weiner et al. 2010; Maierhofer et al. 2014; Deger et al. 2015; Zheng et al. 2018; Takahashi et al. 2022; Pankasem et al. 2024; Xu et al. 2025). In general, plant hormones such as abscisic acid (ABA), jasmonic acid (JA), salicylic acid (SA), as well as reactive oxygen species (ROS), such as superoxide and H2O2, are thought to mediate stomatal closure. In contrast, under certain conditions, ethylene (ET), brassinosteroids (BRs), and other signals (as well as a decrease in the level of stomatal closure-inducing hormones/H2O2) are thought to mediate stomatal opening (Weiner et al. 2010; Hsu et al. 2021; Waadt et al. 2022).

While much is known about the response of stomata to individual stress conditions, very little is known about the response of stomata to conditions of stress combination, and in particular GCFc/MFSC (Zandalinas and Mittler, 2022). For example, under conditions of HS stomata on leaves are open, but under conditions of HS combined with WD (or just WD), stomata on leaves close (Rizhsky et al. 2002, 2004; Sinha et al. 2022, 2023). Recently, it was suggested that the response of stomata to a combination of WD and HS is regulated by an interaction between OPEN STOMATA 1 (OST1) and TARGET OF TEMPERATURE 3 (TOT3; Xu et al. 2025). In addition, it was recently shown that stomata of reproductive tissues of plants respond differently than stomata of leaves, and that under conditions of HS combined with WD stomata on reproductive tissues remain open, while stomata on leaves close, leading to a ‘differential transpiration’ response between reproductive and vegetative tissues that allows the cooling of reproductive tissues during the stress combination (Sinha et al. 2022, 2023; Jensen et al. 2024). Conflicting stomatal responses were further reported for EL stress (stomatal closure) combined with HS (stomatal opening), that resulted in stomata remaining open (Balfagón et al. 2022), and a combination of salt stress (stomatal closing) with HS (stomatal opening), that resulted in stomatal closure or opening depending on the intensity of salt stress conditions (Rivero et al. 2014; Li et al. 2024).

Here, we studied the physiological, genetic and molecular responses of *Arabidopsis thaliana* (L.) Heynh (*A. thaliana*) plants to WD, eCO2, and HS, in all possible combinations, with a focus on stomatal responses. Our findings reveal that plant responses to GCFc are unique and that stomatal regulation under complex conditions of GCFc is mediated by specific sets of regulators/hormones, different from those mediating stomatal regulation under control (CT) or less complex environmental stress conditions. Under the GCFc of WD+HS+eCO2 we found, for example, that stomatal closure requires the function of nitric oxide (NO), OST1 and the S-type anion channel SLAH3, but not the S-type anion channel SLAC1, or other regulators typically found to control stomatal aperture under less complex conditions. Regulation of stomatal aperture under conditions of WD+HS+eCO2, was further found to be different from that under conditions of HS+WD as reported in (Xu et al. 2025). In addition, we report on the hierarchy of stomatal responses in *A. thaliana* plants subjected to WD, W, EL, eCO2, and/or HS, applied in different combinations (*i.e.,* the ‘stomatal hierarchical stress code’).

## 2. MATERIALS AND METHODS

### 2.1 Plant growth and stress treatments

All experiments were conducted in PGC-FLEX growth chambers with flexible control of light, temperature and CO2 levels (Conviron; Winnipeg, Canada). Treatments and plants were randomly assigned to the different chambers between each biological repeat, and wild type (WT) plants were kept at each chamber together with all other plants/treatments/groups to ascertain that responses were the same for all chambers. To control the effects of vapor pressure deficit (VPD) on stomatal aperture (Bauer et al. 2013; Merilo et al. 2018; Grossiord et al. 2020), all chambers were kept at 55% relative humidity for all treatments. Seeds of WT *Arabidopsis thaliana* (L.) Heynh Col-0 and the mutants listed in the Supporting Information (Dataset S1) were obtained from the Arabidopsis Biological Resource Center (ABRC). Seeds of *noa1nia1nia2*, *nia1nia2*, *nox1-1*, and *cue1-5*, mutants were kindly donated by Dr. Óscar Lorenzo (CIALE, Salamanca, Spain). Seeds were germinated and grown on peat pellets (Jiffy-7; Jiffy International, Kristiansand, Norway), under well-watered control conditions of 100 µmol photons s^−1^ m^−2^, 10-h/14-h light/dark cycle at 22 °C/20 °C day/night and 400 μmol mol^−1^ CO2 for 4 to 5 weeks. Whole plants were subjected to the following individual treatments for 1 hour: CT (control; 22 °C, 100 µmol m^-2^ s^-1^, 400 μmol mol^−1^ CO2), EL (elevated light; 22 °C, 1120 µmol photons s^-1^ m^-2^, 400 μmol mol^−1^ CO2), HS (heat stress; 38 °C, 100 µmol m^-2^ s^-1^, 400 μmol mol^−1^ CO2), and/or eCO2 (elevated CO2; 22 °C, 100 µmol m^-2^ s^-1^, 800 μmol mol^−1^ CO2). Wounding (W) and water deficit (WD) were applied under conditions of 22 °C, 100 µmol m^-2^ s^-1^, 400 μmol mol^−1^ CO2. Wounding was applied to half of a single leaf by puncturing with 20 dressmaker pins simultaneously and the non-wounded half leaves were collected after 1 hour for stomatal aperture measurements. For water deficit (WD) treatment, peat pellet weight was monitored until it reached 35 ± 3% of complete peat soil water saturation. To impose the different stress combinations, the individual conditions described above were applied in different combinations. All measurements were conducted between 10AM and noon.

### 2.2 H_2_O_2_ detection

H_2_O_2_ quantification was performed using Amplex-Red (10-acetyl- 3,7-dihydroxyphenoxazine [ADHP]; Thermo Fisher Scientific, Waltham, MA, USA) as described in (Peláez-Vico et al. 2023). Leaves number 6 at the 10-leaf rosette stage were collected from 15 independent plants and immediately frozen. Three independent biological replicates of 5 leaves were ground to fine powder, resuspended in 50 µL 0.1 M TCA (Thermo Fisher Scientific, Waltham, MA, USA), and centrifuged for 15 min at 12,000 × g, 4 °C. The supernatant was buffered with 1 M phosphate buffer pH 7.4, and the pellet was dried and used for dry weight calculation. H2O2 quantification in the supernatant was performed according to the MyQubit-Amplex-Red Peroxide Assay manual (Thermo Fisher Scientific, Waltham, MA, USA), using a calibration curve of H2O2 (Thermo Fisher Scientific, Waltham, MA, USA).

### 2.3 Stomatal aperture and temperature measurements

Stomatal aperture was measured as described in (Peláez-Vico et al. 2023). Briefly, a thin coat of nail polish (450B clear nail protector-Wet N Wild; Markwins Beauty Products, CA, USA) was applied to the abaxial surface of the leaf avoiding the main veins. The nail polish was allowed to dry for approximately 10 min and then peeled off with tweezers. Impressions were mounted facing upward on a microscope slide using double-sided tape (Scotch). An EVOS XL microscope (Invitrogen by Thermo Fisher Scientific, Waltham, MA, USA) at 40X magnification was used to capture stomata images. Images were processed to determine the stomatal pore length and width, measuring the longest axis and the widest point perpendicular at center of the stoma using ImageJ (https://imagej.nih.gov/ij). At least 20 stomata per plant were measured, and a total of 15 plants for each treatment and genotype were included. 20-30 images for each condition and genotype containing around 10-15 stomata on each image were measured (n = 200-500). Stomatal aperture was calculated as the ratio of stomatal pore width to stomatal pore length as described in (Wang et al. 2019). All experiments were conducted between 10 AM and 12 PM. In all experiments, leaf number 6 (one leaf per plant) at the 10-leaf rosette stage was analyzed for all conditions and genotypes. Leaf surface temperature was recorded using a precision infrared thermometer (DX501-RS, Exergen, Watertown, MA, USA) positioned at 1 cm from the leaves. Results include temperature values from at least 20 different plants per treatment.

### 2.4 RNA isolation and RNA-seq analysis

30-day-old WT plants were subjected to eCO2, WD and HS treatments individually and in all possible combinations. Peat pellet weight was monitored until it reached 35 ± 3% of complete peat soil water saturation to induce WD stress alone, and for combinations with eCO2 and HS, the conditions described above were applied for 1 hour. After 1 hour of treatment imposition, leaves (leaf number 6 at 10- leaf rosette stage) were collected and immediately frozen in liquid nitrogen. One leaf per plant from 15 different plants were pooled for each technical repeat and 3 biological repeats were collected for each treatment. RNA was extracted using Plant RNeasy kit (Qiagen, Hilden, Germany) according to the manufacturer instructions. RNA libraries for sequencing were prepared using standard Illumina protocols, and RNA sequencing was performed using NovaSeq 6000 PE150 by Novogene Co. Ltd (https://en.novogene.com/; Sacramento, CA, USA). Processing and expression analysis of RNA-Seq data were performed as described in (Li et al. 2009; Andrews 2010; Trapnell et al. 2012, 2013; Ewels et al. 2016; Kim et al. 2019; Peláez-Vico et al. 2023). Upset plots were generated in upsetr (<u>gehlenborglab.shinyapps.io</u>; Conway et al. 2017). The different stress-, hormone-, and ROS-response transcripts data sets used for comparisons in Figure 1b were obtained from (Peláez-Vico et al. 2023). The transcripts included in “eCO2” dataset were obtained from (Li et al. 2008; Kaplan et al. 2012; Queval et al. 2012; Markelz et al. 2014; Jauregui et al. 2015, 2016; Niu et al. 2016; Higuchi-Takeuchi et al. 2020; Xi et al. 2023).

**FIGURE 1.**
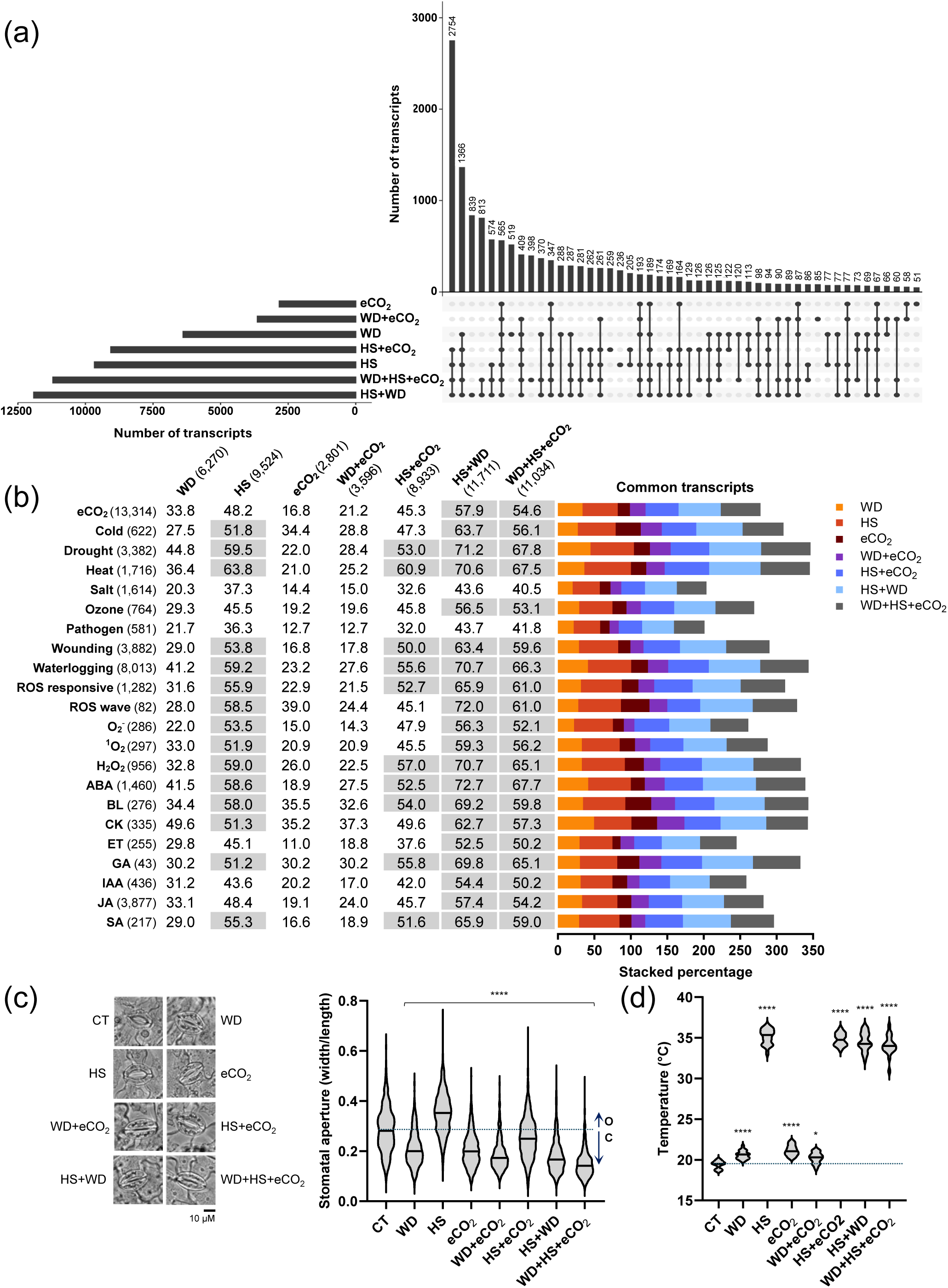
Transcriptomics analysis of *Arabidopsis thaliana* plants subjected to water deficit (WD), heat stress (HS) and elevated CO2 (eCO2) in all combinations. (a) An UpSet plot showing the unique and common transcripts significantly altered in their expression under control conditions (CT), WD, HS, and eCO2 in all possible combinations (See Figure S1a for complete UpSet plot). (b) Representation of different stress, reactive oxygen species (ROS) and hormone response transcripts among the different transcripts significantly altered in their expression under conditions of CT, WD, HS, and eCO2 in all possible combinations. (c) Representative images (left) and graph (right) of stomatal aperture in plants grown under CT conditions, or subjected to WD, HS, and eCO2 in all possible combinations. ‘O’ stands for open, compared to CT, and ‘C’ stands for closed, compared to CT. Measurements of stomatal width and length are shown in Figure S1b. (d) Leaf temperature of plants subjected to CT, WD, HS, and eCO2 in all possible combinations. One-way ANOVA followed by a Dunnett’s multiple comparisons test was used to determine significance. Asterisks denote statistical significance at *p < 0.05, ****p < 0.0001 compared to control. Abbreviations: abscisic acid (ABA), auxin (indole-3-acetic acid; IAA), brassinolide (BL), cytokinin (CK), elevated CO2 (eCO2), ethylene (ET), gibberellic acid (GA), heat stress (HS), jasmonic acid (JA), reactive oxygen species (ROS), salicylic acid (SA), water deficit (WD).

### 2.5 Hormone analysis

Plants were subjected to the different treatments and leaves (leaf number 6 at 10-leaf rosette stage) were collected and immediately frozen in liquid nitrogen. Samples were grounded and freeze-dry for hormone extraction. Hormone analysis was performed as described in (Balfagón et al. 2019) with some modifications. A mixture containing 50 ng of [^2^H6]-ABA, [^13^C]-SA, [^2^H5]-IAA and [^2^H3]-PA and dihydrojasmonic acid was added to around 20-50 mg of grounded, dried leaf tissue. The tissue was homogenized in 2 mL of ultrapure water in a ball mill (MillMix20; Domel, Železniki, Slovenija). After centrifugation at 10,000 × g at 4 °C for 10 min, supernatants were recovered, and pH adjusted to 3 with 30% acetic acid. The water extract was partitioned twice against 2 mL of diethyl ether and the organic layer recovered and evaporated under vacuum in a centrifuge concentrator (Speed Vac; Jouan, Saint Herblain Cedex, France). Then, samples were resuspended in a 90:10 (v/v) H2O:MeOH solution by using a sonicator (Elma S30; Elmasonic, Singen, Germany). After filtering through 0.22 µM polytetrafluoroethylene membrane syringe filters (Albet S.A.; Barcelona, Spain), extracts were directly injected into an ultra-performance UPLC system (Xevo TQ-S; Waters Corp., Milford, MA, USA). Chromatographic separations were performed on a reversed-phase C18 column (Gravity, 50 × 2.1 mm, 1.6 μM particle size; Luna Omega; Phenomenex, Torrance, CA, USA) using a H2O:MeOH (both supplemented with 0.1% formic acid) gradient at a flow rate of 300 μL min^−1^. Hormones were quantified with a triple quadrupole mass spectrometer connected online to the output of the column though an orthogonal Z-spray electrospray ion source. Results were processed using Masslynx v. 4.1 software, and the phytohormone content was quantified with a standard curve prepared with commercial standards as described in (Balfagón et al. 2019).

### 2.6 Data analysis

Statistical analysis was performed using one-way ANOVA followed by Dunnett’s multiple comparisons test, or two-tailed Student’s t-test, in GraphPad. Data distribution is shown as violin plots and the center line marks the median. Asterisks denote statistical significance at *p < 0.05, **p < 0.01, ***p < 0.001, ****p < 0.0001 compared to control.

## 3. RESULTS

### 3.1 Molecular and physiological responses of *A. thaliana* to different combinations of WD, HS, and eCO2

To study the response of *A. thaliana* to GCFc, we focused on the climate change-relevant conditions of HS, WD, and eCO2 and tested their impact on plants in all possible combinations (Figure 1, and S1). Conditions of WD were obtained by gradually decreasing the water content of the planting media, while conditions of HS and eCO2 were applied for 1 hour once plants reached the desired level of WD, simulating a heat wave occurring during conditions of WD under elevated CO2. The rational for using a short-term (1 hour) eCO2 treatment, as opposed to growing plants under constant eCO2 levels, was to determine how stomata will respond to a change in CO2 levels (from ambient to high), and not to address how plants acclimate to stress under constant conditions of eCO2. While plants grown under eCO2 for many generations may have altered metabolism compared to plants subjected to a single generation eCO2 treatment (Yang et al. 2023), several studies have now shown that the genetic and physiological mechanisms that cause stomatal closure in response to a long- or short-term eCO2 treatments are similar (Cheng et al. 1998; Johansson et al. 2020).

To estimate the degree of stress experienced by plants subjected to HS, WD, and eCO2, in all possible combinations, we conducted an integrative transcriptomic, physiological and hormonal analysis of Arabidopsis leaves (Figures 1 and 2; Supporting Information Dataset S2). As shown in Figure 1a and S1a, the combinations of HS+WD and WD+HS+eCO2 resulted in the most comprehensive transcriptomic responses, followed by HS and HS+eCO2, WD, WD+eCO2, and eCO2. In terms of stress responses, HS, HS+eCO2, HS+WD and WD+HS+eCO2 had the highest % representation of significantly expressed transcripts involved in stress/ROS/hormone responses, while eCO2, WD and WD+eCO2 displayed a relatively lower % representation of these transcripts (Figure 1b). Judging by the % representation of the different stress/ROS/hormone transcripts significantly expressed in response to eCO2, this treatment could be considered as an alleviating condition to WD (*i.e.,* WD+eCO2), but not as much to HS, or HS+WD (*i.e.,* HS+eCO2 or WD+HS+eCO2, respectively; Figure 1b). This finding could reflect the conflicting interactions between WD and HS during the HS+WD combination (likely due to opposing stomatal responses; Figure 1c), as well as highlight the fact that a short-term change from ambient to eCO2 may not be as efficient in alleviating the stress conditions experienced by plants during HS+WD stress combination (compare HS+WD with WD+HS+eCO2; Figure 1b).

**FIGURE 2.**
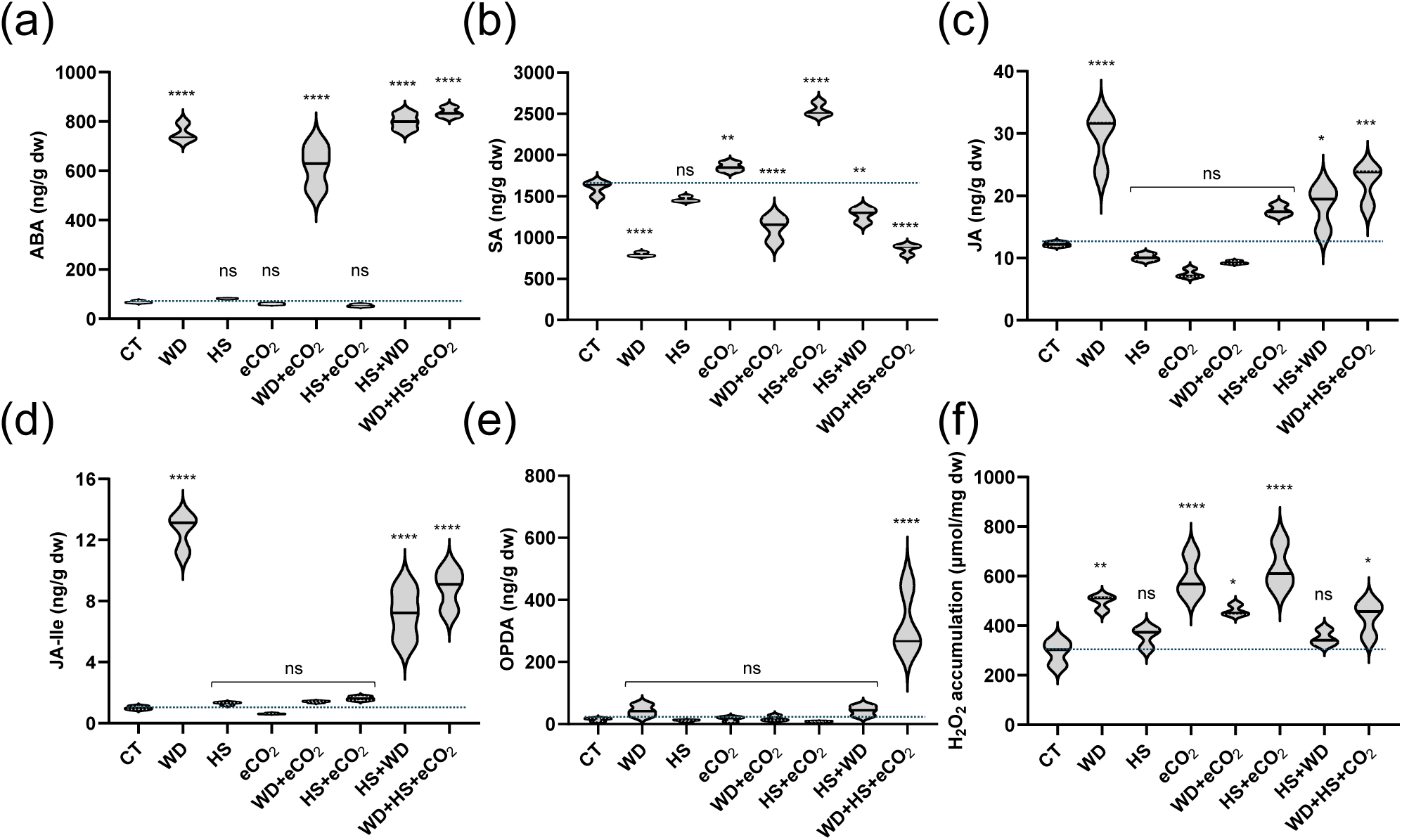
Quantification of hormone and H2O2 levels in plants subjected to different combinations of water deficit (WD), heat stress (HS) and elevated CO2 (eCO2). (a) Abscisic acid (ABA) levels in Arabidopsis plants grown under controlled growth conditions (CT), or subjected to WD, HS, and eCO2 in all possible combinations. (b) Same as in a, but for salicylic acid (SA). (c) Same as in a, but for jasmonic acid (JA). (d) Same as in a, but for JA-Isoleucine (JA-Ile). (e) Same as in a, but for 12-oxo-phytodienoic acid (OPDA). (f) H2O2 accumulation in Arabidopsis plants in response to different combinations of WD, HS and eCO2. One-way ANOVA followed by a Dunnett’s multiple comparisons test was used to determine significance. Asterisks denote statistical significance at *p < 0.05, **p < 0.01, ***p < 0.001, ****p < 0.0001 compared to control. Abbreviations: abscisic acid (ABA), dry weight (dw), elevated CO2 (eCO2), heat stress (HS), hydrogen peroxide (H2O2), jasmonic acid (JA), jasmonic acid-isoleucine (JA-Ile), non-significance (ns), 12-oxo- phytodienoic acid (OPDA), salicylic acid (SA), water deficit (WD).

Measurements of stomatal aperture 1 hour following stress/treatment application revealed that in response to WD and eCO2 stomatal aperture decreased, while in response to HS stomatal aperture increased (Figure 1c and S1b). The effects of the different individual treatments on stomatal aperture, measured in our study, therefore agreed with previous publications (*e.g.,* Rizhsky et al. 2004; Johansson et al. 2020; Sinha et al. 2022). Synergistic (WD+eCO2) and antagonistic (HS+WD and HS+eCO2) effects were identified between the different individual stresses (Figure 1c; the antagonistic effect between WD and HS was originally discovered by Rizhsky et al. 2002, 2004). In response to the WD+HS+CO2 combination, stomata were closed, revealing that under this GCFc, like the results for soybean plants subjected to MFSC (Peláez-Vico et al. 2024a), the overall response of plants is to close their stomata. Measurements of leaf temperature, that generally reflect stomatal responses (due to the cooling effect of transpiration that requires stomatal opening; Rizhsky et al. 2002, Sinha et al. 2022), supported our stomatal aperture measurements (Figure 1d).

### 3.2 Hormonal responses of *A. thaliana* to the different combinations of WD, HS, and eCO_2_

Leaf tissues sampled in parallel to the transcriptomics and stomatal analyses shown in Figure 1, was further subjected to total leaf hormonal and H2O2 content analyses (Figure 2). The levels of ABA and its catabolite phaseic acid (PA) were high in plants subjected to WD and any growth condition that included WD (Figures 2a and S2; in agreement with the expression of ABA-response transcripts; Figure 1b), correlating with stomatal closure (Figure 1c). In contrast, SA levels displayed an opposite correlation with WD (Figure 2b). The levels of JA and JA-Isoleucine (JA-Ile) were high in plants subjected to WD, but suppressed in plants grown under eCO2, unless all 3 growth conditions were combined (Figure 2c, 2d). In contrast, 12-oxophytodienoic acid (OPDA) levels specifically increase in plants subjected to the combination of all 3 growth factors (Figure 2e). In contrast to ABA, total leaf H2O2 levels did not correspond with stomatal aperture closure and revealed an interesting positive relationship with eCO2 (Figure 2f). Thus, while plants grown under all experimental growth conditions contained higher H2O2 levels compared to control, plants grown under WD, eCO2, and any combination that included eCO2, displayed higher H2O2 levels. Interestingly, total auxin (indole-3-acetic acid; IAA) levels were suppressed under all growth conditions, besides the 3-growth conditions combination (Figure S3). In contrast to ABA, the pattern of JA-, SA-, and H2O2-response transcripts did not follow the accumulation pattern of their corresponding compounds; Figures 1, 2).

To further determine the role of hormones/H2O2 in regulating stomatal responses under GCFc, we studied stomatal responses and hormone levels in different mutants impaired in ABA, JA, and ROS signaling (Figures 3 and S4). The *aba2-11* mutant, deficient in ABA biosynthesis, was impaired in stomatal closure in response to all growth conditions, except HS+WD, as well as impaired in stomatal opening during HS (Figures 3a and S4). As expected, ABA (and PA) levels were low in the *aba2-11* mutant (Figure S5). In contrast, OPDA levels were high in the 3-growth conditions combination in the *aba2-11* mutant (Figure S5), suggesting that OPDA accumulation alone (Figure 2e) is insufficient to signal stomatal closure under these conditions. In addition, JA-Ile levels were similar to those of WT in the *aba2-11* mutant (Figures 2d and S5). Taken together, these results suggest that total leaf JA-Ile or OPDA accumulation are not associated with most stomatal responses of plants to the different stresses/growth conditions, or their combination. In a mutant impaired in JA perception (*coi1*), stomatal opening during HS and stomatal closure in response to eCO2, or HS+eCO2 were less prominent (Figures 3b and S4). In contrast, stomatal closure associated with WD or WD in combination with any other condition was not completely eliminated (Figures 3b and S4). These results suggest that JA sensing is required for stomatal opening during HS and stomatal closing by eCO2 but is not required for stomatal closure by WD and many of the stress/growth condition combinations. ABA and PA levels were mostly similar to WT (besides HS+WD) in the *coi1* mutant, and JA-Ile and OPDA levels were lower than WT under the 3-growth conditions combination (Figure S5). Stomatal responses of the JA biosynthesis mutants *aos1* and *opr3* were mostly similar to WT, with the exception that stomatal opening in response to HS and stomatal closure in response to HS+eCO2 were suppressed (somewhat similar to the *coi1* mutant; Figures 3c, 3d, and S4). Compared to WT, JA and JA-Ile levels were suppressed in the *opr3* mutant under all stress conditions (Figure S5). While OPDA levels were higher than control in the *opr3* mutant, they were not as high as in WT in response to the 3-growth factor combination (Figure S5). As with the *coi1* mutant, the pattern of ABA and PA accumulation was similar to that of WT (albeit lower) in the *opr3* mutant (Figure S5). The stomatal responses of the *rbohD* mutant were similar to those of WT, albeit suppressed in their intensity under most growth conditions/combinations (Figures 3e and S4). In *NahG* plants, that actively degrade SA, stomatal aperture in responses to WD, HS, and WD+eCO2 were not significantly altered compared to WT, while stomatal aperture response to HS+WD, eCO2, or any other combination that included eCO2 (including WD+HS+eCO2) were suppressed (Figures 3f and S4).

**FIGURE 3.**
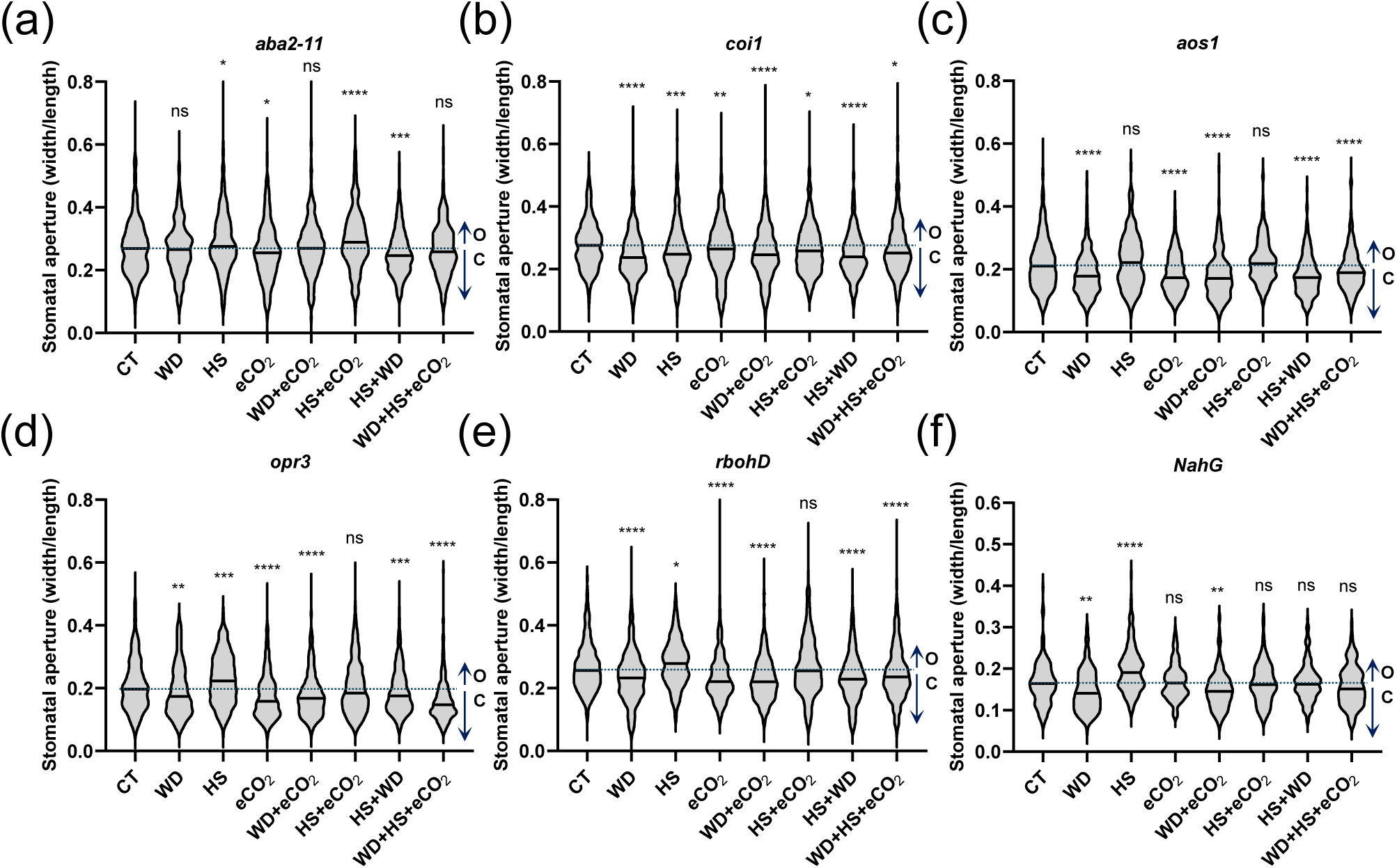
Stomatal aperture of *Arabidopsis thaliana* mutants deficient in the biosynthesis or sensing of different plant hormones and reactive oxygen species (ROS), under conditions of water deficit (WD), heat stress (HS) and/or elevated CO2 (eCO2). Stomatal aperture of *Arabidopsis thaliana* mutants deficient in the levels of abscisic acid (ABA), *aba2-11* (a), jasmonic acid (JA), *coi1* and *aos1* (b and c), 12-oxo-phytodienoic acid (OPDA), *opr3* (d), ROS, *rbohD* (e), and salicylic acid (SA), *NahG* (f), grown under controlled growth conditions (CT), or subjected to WD, HS, and eCO2 in all possible combinations. ‘O’ stands for open, compared to CT, and ‘C’ stands for closed, compared to CT. Measurements of stomatal width and length are shown in Figure S4. Details of the different mutants are given in the Supporting Information (Dataset S1). One-way ANOVA followed by a Dunnett’s multiple comparisons test was used to determine significance. Asterisks denote statistical significance at *p < 0.05, **p < 0.01, ***p < 0.001, ****p < 0.0001 compared to control. Details of the different mutants are given in the Supporting Information (Dataset S1). Abbreviations: abscisic acid (aba), allene oxide synthase (aos), coronatine-insensitive (coi), elevated CO2 (eCO2), heat stress (HS), non-significance (ns), oxylipin 12-oxo-phytodienoic acid reductase (opr), respiratory burst oxidase homolog (rboh), salicylate hydroxylase (NahG), water deficit (WD).

Taken together, the results shown in Figures 2, 3, S4, and S5, suggest that stomatal closure responses under all conditions (except HS+WD) could primarily be mediated by ABA, while stomatal opening responses during HS and stomatal closure responses under conditions of eCO2, might require ABA, JA, and SA. In addition, the intensity (but not direction) of all stomatal responses appears to be enhanced by RBOHD-produced ROS. The only plant hormone that appeared to be involved in stomatal responses to HS+WD and WD+HS+eCO2 was SA (with ABA also required for stomatal closure by WD+HS+eCO2, but not HS+WD).

### 3.3 Downstream mechanisms mediating stomatal responses to the different growth conditions and their combination

To further determine the molecular mechanisms that regulate stomatal responses under conditions of HS, WD, and/or eCO2, and to link these mechanisms with changes in hormonal levels, we studied stomatal responses of selected mutants impaired in different channels and kinases known to regulate stomatal aperture (Figure 4a; Waadt et al. 2022; Jezek and Blatt, 2017; Blatt, 2024). As shown in Figures 4b and S6 OST1, a central and multifaceted regulator of stomatal responses (Belin et al. 2006; Deng et al. 2021), was required for stomatal responses (open or close) to all different growth conditions and their combinations, placing it at a key junction for stomatal regulation under stress combination. In contrast to OST1, AHA1 was not required for stomatal opening under conditions of HS, or for stomatal closure under conditions of WD+HS+eCO2. AHA1 was, however, required for stomatal closure under conditions of WD, eCO2 and all possible two-condition combinations, including HS+WD (Figures 4c and S6). In agreement with (Xu et al. 2025), TOT3 was required for stomatal opening during HS. TOT3 was also required for stomatal closure under WD, but not for any other treatment/combination, including HS+WD (Figures 4d and S6). In contrast to OST1, AHA1 and TOT3, the receptor-like kinase (LRR- RLK) GUARD CELL HYDROGEN PEROXIDE-RESISTANT1 (GHR1) was only required for stomatal opening during HS and stomatal closure during a combination of WD+eCO2 (Figures 4e and S6). Taken together, the findings presented in Figures 4 and S6 reveal that OST1 has a central role in regulating stomatal responses under multiple growth conditions, including the combination of WD+HS+eCO2. In contrast, AHA1, TOT3, or GHR1 that were involved in responses to HS, WD, and/or eCO2, were not required for stomatal closure under conditions of WD+HS+eCO2. These findings highlight the unique mode of regulation of stomatal aperture under complex conditions of growth conditions.

**FIGURE 4.**
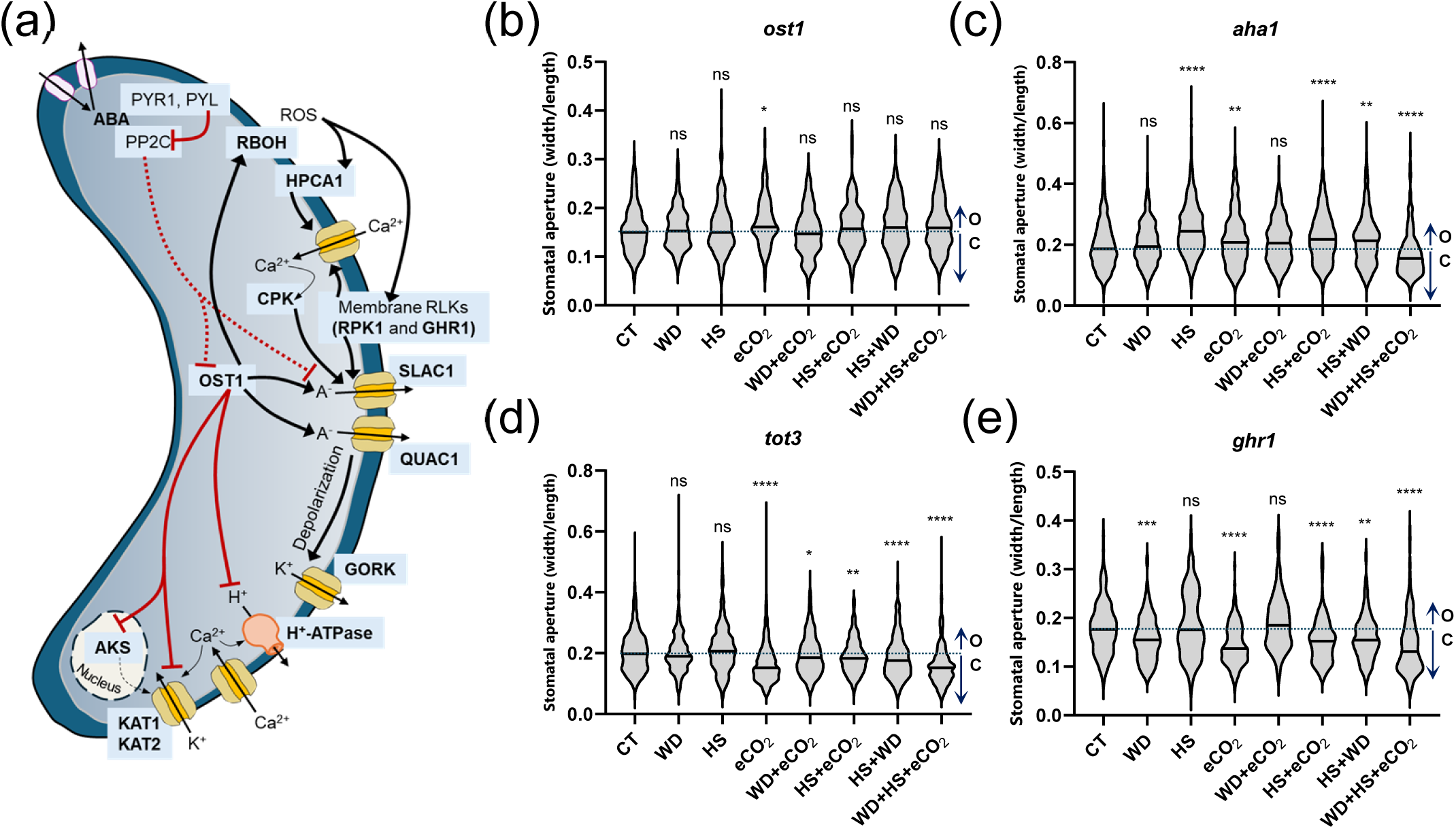
OPEN STOMATA 1 is required for stomatal closure in plants subjected to a combination of water deficit (WD), heat stress (HS) and elevated CO2 (eCO2). (a) A model showing some of the key players in stomatal aperture regulation in plants. Adapted from (Jezek and Blatt 2017; Waadt et al. 2022; Jaslan et al. 2023; Blatt, 2024). (b) Stomatal aperture of *Arabidopsis thaliana* mutants deficient in OPEN STOMATA 1 (*ost1*), grown under controlled growth conditions (CT), or subjected to WD, HS, and eCO2 conditions in all possible combinations. (c) Same as in b, but for mutants deficient in the plasma membrane H^+^-ATPase AHA1 (*aha1*). (d) Same as in b, but for mutants deficient in the TARGET OF TEMPERATURE 3 (TOT3) kinase (*tot3*). (e) Same as in b, but for mutants deficient in the GUARD CELL HYDROGEN PEROXIDE-RESISTANT1 (GHR1) protein (*ghr1*). ‘O’ stands for open, compared to CT, and ‘C’ stands for closed, compared to CT. Details of the different mutants are given in the Supporting Information (Dataset S1) and results for additional alleles are shown in Figure S8. Measurements of stomatal width and length are shown in Figure S6. One-way ANOVA followed by a Dunnett’s multiple comparisons test was used to determine significance. Asterisks denote statistical significance at *p < 0.05, **p < 0.01, ***p < 0.001, ****p < 0.0001 compared to control. Abbreviations: abscisic acid (ABA), ABA-RESPONSIVE KINASE SUBSTRATE (AKS), CA^2+^-DEPENDENT PROTEIN KINASE (CPK), elevated CO2 (eCO2), GUARD CELL HYDROGEN PEROXIDE-RESISTANT1 (GHR1), GUARD CELL OUTWARD RECTIFYING K+ CHANNEL (GORK), HYDROGEN-PEROXIDE-INDUCED Ca^2+^ INCREASES 1 (HPCA1), heat stress (HS), CHANNEL IN ARABIDOPSIS THALIANA 1 (KAT1), CHANNEL IN ARABIDOPSIS THALIANA 2 (KAT2), non-significance (ns), OPEN STOMATA 1 (OST1), PROTEIN PHOSPHATASE TYPE 2C (PP2C), PYRABACTIN RESISTANCE 1-LIKE (PYL), PYRABACTIN RESISTANCE 1 (PYR1), QUICK-ACTIVATING ANION CHANNEL 1 (QUAC1), RESPIRATORY BURST OXIDASE HOMOLOG (RBOH), RECEPTOR-LIKE KINASES (RLKs), reactive oxygen species (ROS), RECEPTOR-LIKE PROTEIN KINASE1 (RPK1), SLOW ANION CHANNEL-ASSOCIATED 1 (SLAC1), water deficit (WD).

### 3.4 SLAH3 and NO are required for stomatal closure during a combination of WD+HS+eCO_2_

To determine the molecular mechanisms involved in stomatal closure under complex growth conditions, we focused on the combination of WD+HS+eCO2 and screened additional mutants impaired in different channels and kinases, as well as hormone biosynthesis, involved in the regulation of stomatal aperture (Figures 5, S7 and S8). As shown in Figure 5a, stomatal closure was suppressed in mutants deficient in NO biosynthesis (*nia1nia2* and *noa1nia1nia2*), but not in mutants that over-accumulate NO (*nox1* and *cue1*). In addition, as shown in Figures 5b, S7 and S8, among the different anion channel mutants deficient in stomatal regulation, only SLAH3, and not SLAC1 that is typically involved in stomatal closure responses mediated by OST1 (Jezek and Blatt, 2017; Jaślan et al. 2023; Blatt, 2024), was deficient in stomatal closure under conditions of WD+HS+eCO2. Taken together, the findings presented in Figures 2-5, and S4-S8 reveal that stomatal closure under conditions of WD+HS+eCO2 required SA, ABA, NO, OST1 and SLAH3, constituting a potential new module for stomatal regulation under complex growth/stress conditions (Figure 5d).

**FIGURE 5.**
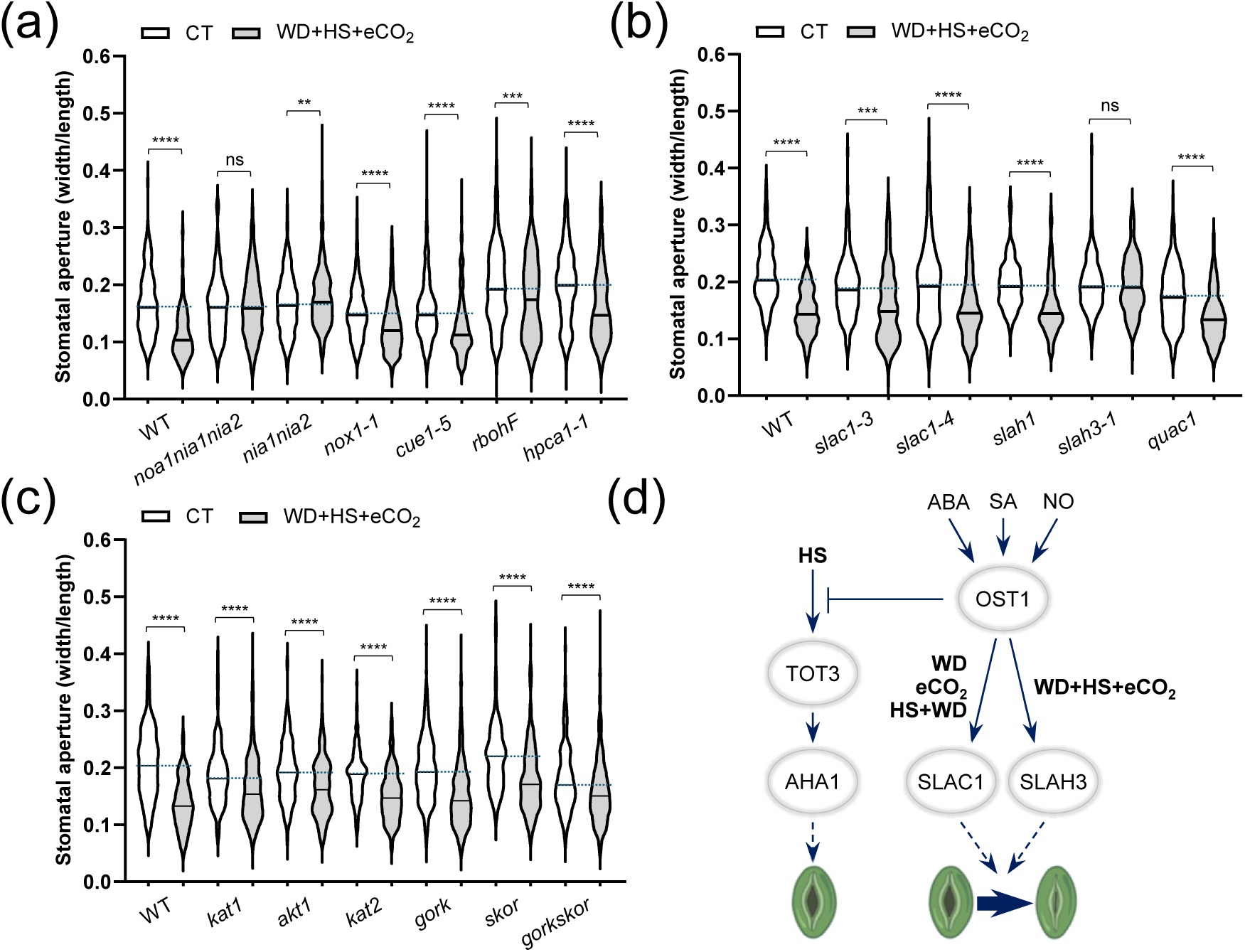
Nitric oxide signaling and SLAH3 are required for stomatal closure in plants subjected to a combination of water deficit (WT), heat stress (HS) and elevated CO2 (eCO2), and a model. (a) Stomatal aperture of *Arabidopsis thaliana* mutants deficient in nitric oxide (NO) and reactive oxygen species (ROS) signaling, grown under control conditions (CT), or subjected to a combination of WD, HS, eCO2. (b) Same as in a, but for *A. thaliana* mutants deficient in the function of selected anion channels involved in stomatal aperture regulation. (c) Same as in a, but for *A. thaliana* mutants deficient in the function of selected cation channels involved in stomatal aperture regulation. (d) A model for stomatal aperture regulation in plants subjected to a combination of WD, HS, and eCO2. OST1 and SLAH3 are shown to be required for stomatal closure under a combination of WD+HS+eCO2. The model was drawn based on the results obtained by the current study, the results of (Xu et al. 2025), as well as results summarized by (Weiner et al. 2010; Kollist et al. 2014; Takahashi et al. 2022; Waadt et al. 2022; Melotto et al. 2024). Details of the different mutants are given in the Supporting Information (Dataset S1) and results for additional alleles are shown in Figure S8. Measurements of stomatal width and length are shown in Figure S7. Two-tailed Student’s t-test was used to determine significance. Asterisks denote statistical significance at **p < 0.01, ***p < 0.001, ****p < 0.0001. Abbreviations: abscisic acid (ABA), ARABIDOPSIS H^+^-ATPase 1 (AHA1), K^+^ transporter 1 (akt1), chlorophyll a/b binding protein underexpressed 1 (cue), elevated CO2 (eCO2), gated outwardly-rectifying K^+^ channel (gork), hydrogen-peroxide-induced Ca^2+^ increases 1 (HPCA1), heat stress (HS), channel in *Arabidopsis thaliana* 1 (kat1), channel in *Arabidopsis thaliana* 2 (kat2), nitrate reductase (nia), nitric oxide (NO), nitric oxide associated 1 (noa), nitrous oxide overexpressor 1 (nox1), non-significance (ns), OPEN STOMATA 1 (OST1), quick-activating anion channel 1 (quac1), respiratory burst oxidase homolog (rboh), salicylic acid (SA), stelar K^+^ outward rectifier (skor), SLOW ANION CHANNEL-ASSOCIATED 1 (SLAC1), SLAC1 homologue 1 (slah1), SLAC1 homologue 3 (SLAH3), TARGET OF TEMPERATURE 3 (TOT3), water deficit (WD).

### 3.5 Hierarchy in stomatal responses to stress/growth condition combination

To further determine the hierarchy of stomatal responses to different stresses/growth conditions that could be involved in GCFc/MFSC/stress combination, we subjected *A. thaliana* plants to CT, WD, and/or a short-term (1 hour) treatment of EL, W, eCO2, and/or HS, in different combinations, and measured their stomatal aperture 1 hour following stress/treatment application (Figures 6a and S9). Based on the degree of stomatal aperture and the overall response to each combination (open or close), we then established what stress/growth condition had priority in determining stomatal aperture during all the different two-stress/condition combinations, and from this analysis defined the ‘stomatal hierarchical code’ (Figure 6b). Under the conditions we used, WD had the highest priority over all stresses/growth conditions in determining stomatal aperture, and all other stresses/growth conditions had a higher priority over HS (Figure 6).

**FIGURE 6.**
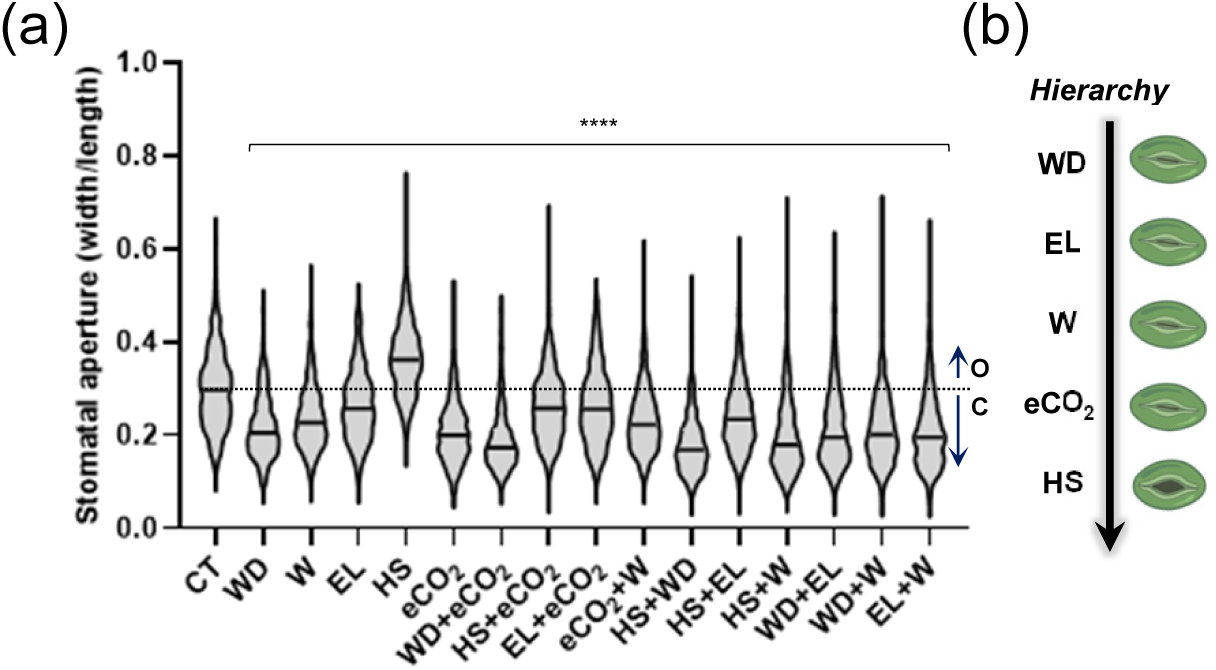
Hierarchy in stomatal stress regulation during stress combination in the flowering plant *Arabidopsis thaliana*. (a) Stomatal aperture of plants grown under controlled growth conditions (CT), or subjected to water deficit (WD), wounding (W), excess light (EL), heat stress (HS), and/or elevated CO2 (eCO2) in different combinations. ‘O’ stands for open, compared to CT, and ‘C’ stands for closed, compared to CT. (b) A model for the hierarchy of stomatal stress regulation during different stress combinations. Measurements of stomatal width and length are shown in Fig. S9. One-way ANOVA followed by a Dunnett’s multiple comparisons test was used to determine significance. Asterisks denote statistical significance at ****p < 0.0001 compared to control. Abbreviations: elevated CO2 (eCO2), excess light (EL), heat stress (HS), water deficit (WD), wounding (W).

## 3. DISCUSSION

As conditions on our planet are becoming more complex and extreme due to global warming, climate change, and increased pollution, a new focus is needed on studying how plants respond to conditions of GCFc/MFSC (Zandalinas et al. 2021; Mittler et al. 2022; Zandalinas and Mittler 2022; Sato et al. 2024). Addressing how conditions of GCFc impact the molecular and physiological responses of *A. thaliana*, our work reveals that: 1. The transcriptomics and hormonal responses of plants to conditions of GCFc are unique and cannot be extrapolated from simply studying the response of plants to each individual stress involved in the GCFc (Figures 1, 2); 2. Under conditions of GCFc stomatal aperture is regulated by a blend of unique and overlapping pathways that are different from those operating under less complex stress conditions (Figures 3-5); and 3. Stomatal responses follow a hierarchical code that is based on the type of each stress/growth condition involved in a combination of two or more stresses/conditions simultaneously impacting a plant (Figures 1c, 6). Focusing on stomatal responses to WD+HS+eCO2, we further identified a potential new regulatory module that involves OST1 and SLAH3, and requires the function of SA, NO, and ABA, to induce stomatal closure under the combination of WD+HS+eCO2 (Figure 5d).

Although the conflicting effects between WD and HS on stomatal aperture regulation, occurring during a WD+HS combination, were discovered over 20 years ago (Rizhsky et al. 2002, 2004), the detailed mechanisms controlling this response were only recently uncovered (Xu et al. 2025). Here, we reveal, however, that the mechanisms regulating stomatal responses under conditions of WD+HS+eCO2 are different from those controlling stomatal responses to HS+WD (Figures 3-5). This finding highlights the complexity of stomatal regulation under conditions of stress combination and suggests that more studies are needed to untangle the ‘stomatal hierarchical code’ of plants (Figures 5d, 6). Moreover, as stomatal responses of vegetative and reproductive tissues of different crops are different from each other under conditions of stress combination (Sinha et al. 2022, 2023; Jensen et al. 2024), studies defining the molecular mechanisms controlling stomatal responses to compound growth conditions should focus on both vegetative and reproductive tissues.

When addressing the potential of eCO2 to alleviate or worsen certain stress/stress combination conditions, revealed in the current study by measuring leaf temperature (Figure 1d) and transcriptional responses (Figures 1a, 1b, S1a), it is important to remember that in our study we used a short-term (1 hour) treatment of eCO2. In contrast, other studies exposed to, or grew plants, over long periods of time under conditions of eCO2 (Cheng et al. 1998; Johansson et al. 2020; Yang et al. 2023). Although in general our transcriptomics analysis supports previous studies revealing that eCO2 alleviates some stress conditions in plants (Figures 1, S1a; Leakey et al. 2009; Thomey et al. 2019), and prior studies showed similarity between the genetic mechanisms that control stomatal closure under long- and short-term eCO2 exposure (Johansson et al. 2020), care should be exercised when interpreting our short-term eCO2 results and their relevance to plants grown under constant conditions of eCO2.

The stomatal hierarchy of stress/growth condition combination revealed by our work (Figure 6b) should be studied in other plant species, as well as under additional/different stress/stress combination conditions, applied in different intensities, order of application, and complexity, as well as over different timescales. This is exemplified by our findings for HS, EL, and HS+EL (Figure 6a). While stomatal responses to the individual conditions of HS and EL recorded in the current work are in agreement with previous studies, the outcome of the EL+HS combination reported here (Figure 6a) is different from that reported by previous studies (Balfagón et al. 2019; Zandalinas et al. 2020), albeit in (Zandalinas et al. 2020) the systemic (but not local) stomatal response of plants to the HS+EL combination agrees with the current study. Our findings that stomatal opening in response to HS occurred in the *aha1* mutant (Figure 4c; conflicting with the model proposed in Xu et al. 2025), are a further indication that more studies are needed to untangle stomatal regulation under conditions of stress combination. Depending on different environmental parameters, stomatal aperture responses to HS could therefore be mediated by more than one mechanism (Pankasem et al. 2024; Xu et al. 2025). The hierarchy in stomatal responses and the conflicting interactions between different stresses/growth conditions combinations likely depend on different factors such as stress intensity, duration, order of stresses applied, and VPD, and can be different in different plants grown under different experimental conditions (Blatt et al. 2022; Zandalinas and Mittler 2022). Stomatal responses to GCFc should therefore be studied in more detail in multiple plants and crops under multiple conditions (as well as in vegetative and reproductive tissues; Sinha et al. 2022, 2023; Jensen et al. 2024).

The interactions between the different plant hormones and the mechanisms they control in plants during stress combination should also be studied in more detail. For example, not much is known about how OST1 regulates SLAH3. SLAH3 was shown to be regulated by calcium signaling via calcineurin B-like proteins (CBLs) and calcium-dependent protein kinases (CPKs), but so was SLAC1 (Belin et al. 2006; Kollist et al. 2014; Maierhofer et al. 2014; Deger et al. 2015; Jezek and Blatt, 2017; Zheng et al. 2018; Deng et al. 2021; Blatt et al. 2022; Jaślan et al. 2023; Blatt, 2024). It is therefore unclear whether OST1 controls SLAH3 by direct phosphorylation, or through calcium signaling. Further studies are needed to resolve this question. Likewise, compared to ABA, much less is known about how SA and NO control stomatal responses during stress (Hsu et al. 2021; Waadt et al. 2022). Additional studies are therefore needed to address the interactions between NO, SA, and ABA during stress combination, and these studies should be conducted using whole leaves and reproductive tissues, as well as single-cell methods focusing on guard cells.

Taken together, our work reveals a high level of complexity in the molecular, hormonal, and stomatal responses of plants to conditions of GCFc. This complexity should be taken into consideration when attempting to develop crops with heightened resilience to climate change, as well as when considering the effects of GCFc on different ecosystems worldwide. As conflicting stomatal responses could cause higher plant temperatures (due to reduced transpiration) and suppressed photosynthetic activity, resulting in decreased survival, growth, and yield during heat waves or heat waves combined with other stresses such as drought, waterlogging, or high ozone levels (Rizhsky et al. 2002, 2004; Zandalinas et al. 2021; Sinha et al. 2022; Zandalinas and Mittler, 2022; Sinha et al. 2023), we need to understand how stomatal responses are controlled under complex environmental conditions. Such studies conducted in different plants and under different conditions are crucial to our understanding of how GCFc impact different ecosystems and agro- ecosystems worldwide. We hope that our study will facilitate such essential investigations as well as draw attention to their importance.

## Acknowledgments

This work was supported by funding from the National Science Foundation (IOS-2414183; IOS-2110017, IOS-2343815) and the Interdisciplinary Plant Group, and University of Missouri, Columbia.

## Author Contributions

M.A.P.V., S.I.Z., M.F.L.C. and R.S. performed experiments and analyzed the data. A.G. and T.J. analyzed the transcriptomics data. M.A.P.V., F.B.F., and R.M. designed experiments. M.A.P.V., F.B.F., R.S., S.I.Z., and R.M wrote the manuscript and/or provided feedback. R.M. and F.B.F. provided financial support.

## Competing Interest Statement

The authors declare no competing interests.

## Data availability

The data that supports the findings of this study are available in the text, figure, and supporting information of this article. RNA-seq data files are available at GEO (GSE284452).

## Supporting Information

Dataset S1 Details of the different mutants used for the study.

Dataset S2 Transcripts significantly altered in wild type Arabidopsis thaliana plants in response to the different stress/growth conditions.

Figure S1 A complete UpSet plot showing the unique and common transcripts significantly altered in their expression under conditions of CT, WD, HS, and eCO2 in all possible combinations (In support of Figure 1a).

Figure S2 Phaseic acid (PA) levels in plants grown under controlled growth conditions (CT), or subjected to water deficit (WD), heat stress (HS), and elevated CO2 (eCO2) in all possible combinations (In support of Figure 2).

Figure S3 Auxin (IAA) levels in plants grown under controlled growth conditions (CT), or subjected to water deficit (WD), heat stress (HS), and elevated CO2 (eCO2) in all possible combinations (In support of Figure 2).

Figure S4 Measurements of stomatal width and length for the results shown in Figure 3. In support of Figure 3.

Figure S5 Hormone levels in aba2-11, coi1, and opr3 mutants grown under controlled growth conditions (CT), or subjected to water deficit (WD), heat stress (HS), and elevated CO2 (eCO2) in all possible combinations (In support of Figure 2).

Figure S6 Measurements of stomatal width and length for the results shown in Figure 4. In support of Figure 4.

Figure S7 Measurements of stomatal width and length for the results shown in Figure 5. In support of Figure 5.

Figure S8 Stomatal aperture of different mutants (second alleles) grown under controlled growth conditions (CT), or subjected to water deficit (WD), heat stress (HS), and elevated CO2 (eCO2) in different combinations (In support of Figures 4, 5).

Figure S9 Measurements of stomatal width and length for the results shown in Figure 6a. In support of Figure 6.

---

**Figure S1.**
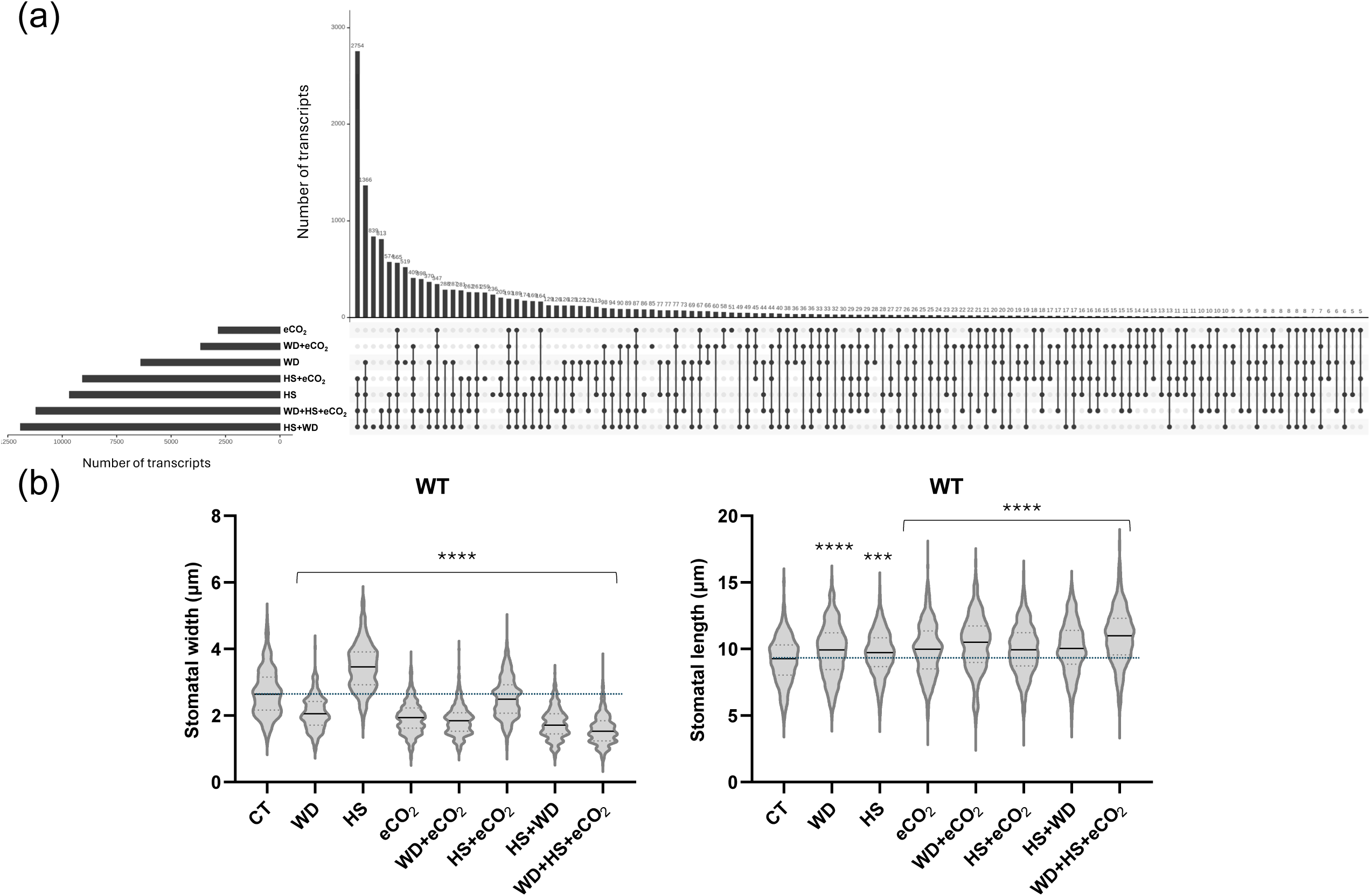
A complete UpSet plot showing the unique and common transcripts significantly altered in their expression under conditions of CT, WD, HS, and eCO2 in all possible combinations (In support of Figure 1a). Abbreviations: elevated CO2 (eCO2), heat stress (HS), water deficit (WD). Measurements of stomatal width and length for the results shown in Figure 1c. In support of Figure 1.

**Figure S2.**
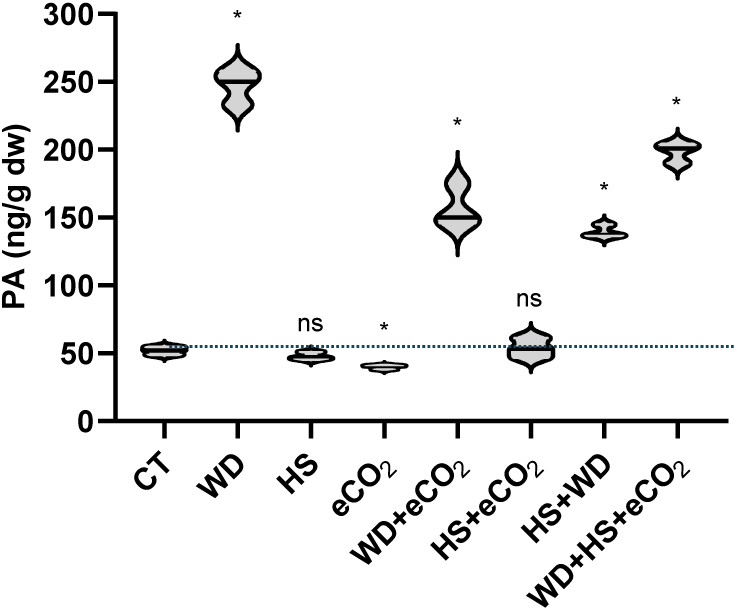
Phaseic acid (PA) levels in plants grown under controlled growth conditions (CT), or subjected to water deficit (WD), heat stress (HS), and elevated CO_2_ (eCO_2_) in all possible combinations (In support of Figure 2). Two-tailed Student’s t-test was used to determine significance. Asterisks denote statistical significance at *p < 0.05 compared to control. Abbreviations: control (CT), elevated CO_2_ (eCO_2_), heat stress (HS), non-significance (ns), phosphatidic acid (PA), water deficit (WD).

**Figure S3.**
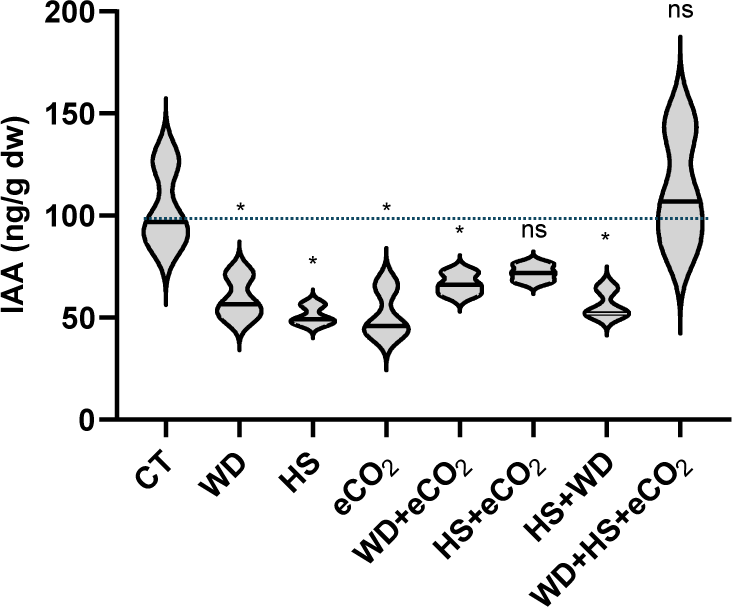
Auxin (IAA) levels in plants grown under controlled growth conditions (CT), or subjected to water deficit (WD), heat stress (HS), and elevated CO_2_ (eCO_2_) in all possible combinations (In support of Figure 2). Two-tailed Student’s t-test was used to determine significance. Asterisks denote statistical significance at *p < 0.05 compared to control. Abbreviations: control (CT), elevated CO_2_ (eCO_2_), heat stress (HS), indole-3-acetic acid (IAA), non-significance (ns), water deficit (WD).

**Figure S4.**
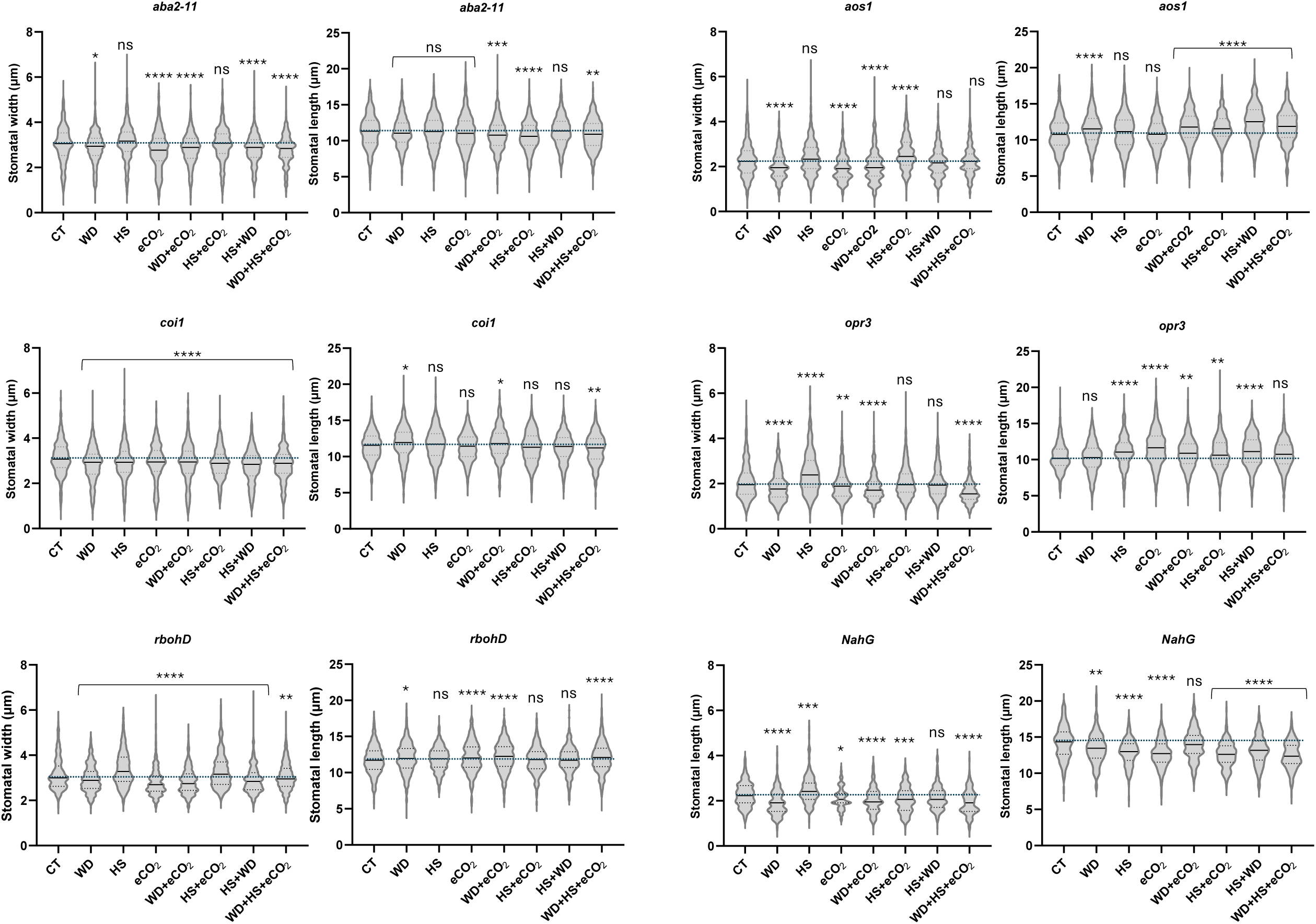
Measurements of stomatal width and length for the results shown in Figure 3. In support of Figure 3.

**Figure S5.**
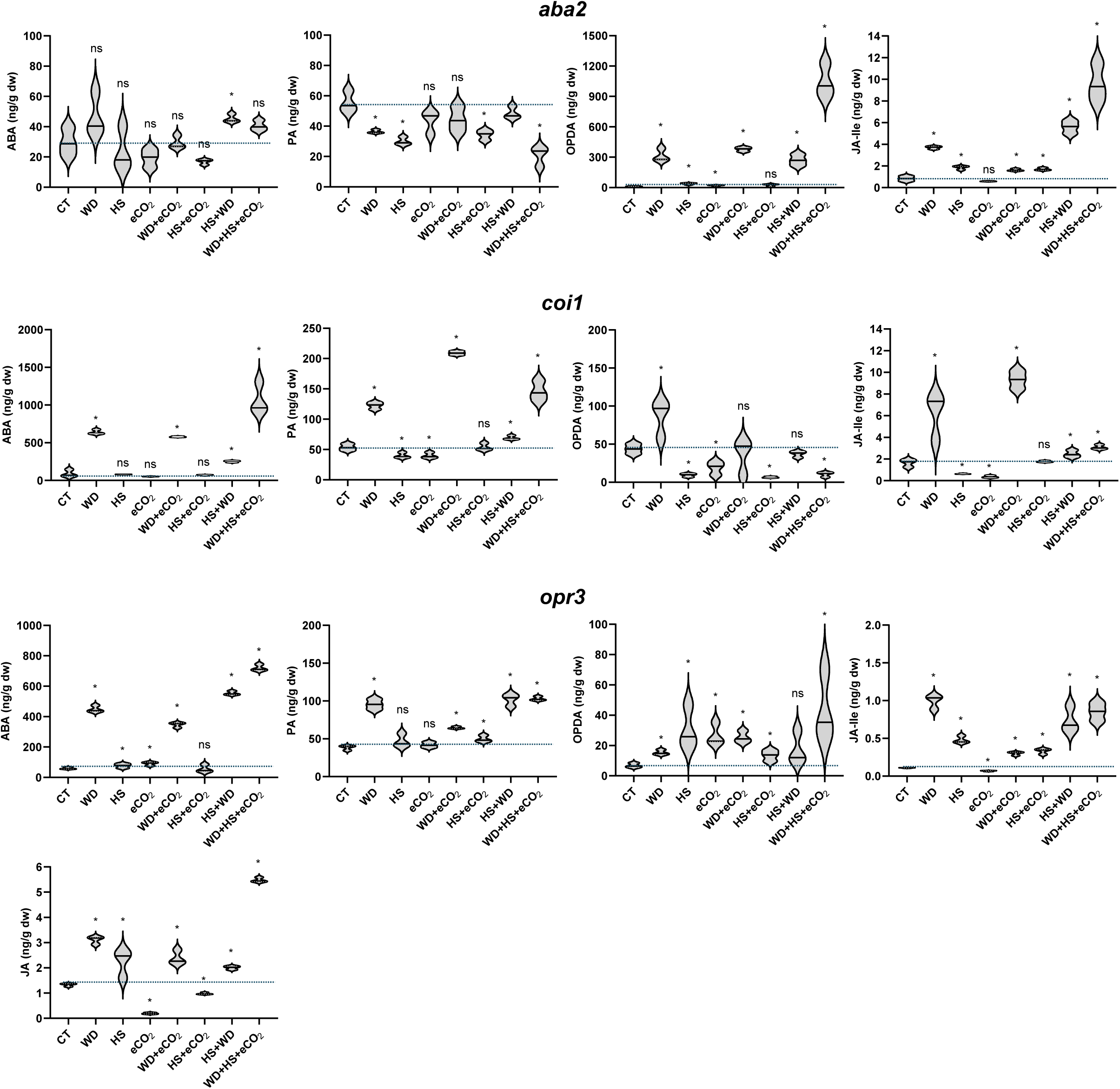
Hormone levels in aba2-11, coi1, and opr3 mutants grown under controlled growth conditions (CT), or subjected to water deficit (WD), heat stress (HS), and elevated CO2 (eCO2) in all possible combinations (In support of Figure 2). Two-tailed Student’s t-test was used to determine significance. Asterisks denote statistical significance at *p < 0.05 compared to control. Details of the different mutants are given in the Supporting Information (Dataset S1). Abbreviations: abscisic acid (ABA), coronatine-insensitive (coi), control (CT), elevated CO2 (eCO2), heat stress (HS), Jasmonic Acid (JA), Jasmonic Acid-Isoleucine (JA-Ile), 12-oxophytodienoic acid (OPDA), oxylipin 12-oxophytodienoic acid (OPDA), oxylipin 12-oxophytodienoic acid reductase (opr), phosphatidic acid (PA), water deficit (WD).

**Figure S6.**
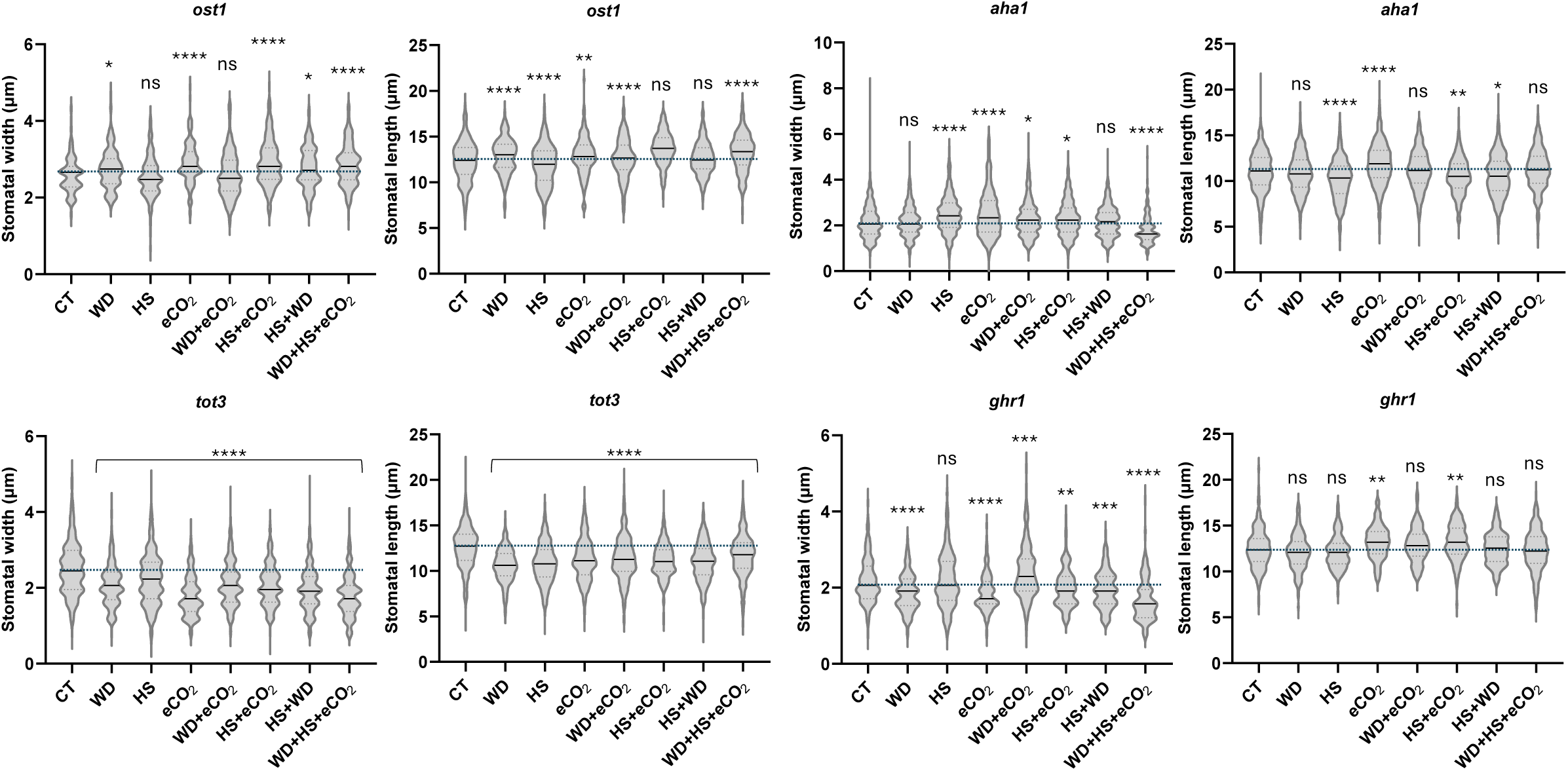
Measurements of stomatal width and length for the results shown in Figure 4. In support of Figure 4.

**Figure S7.**
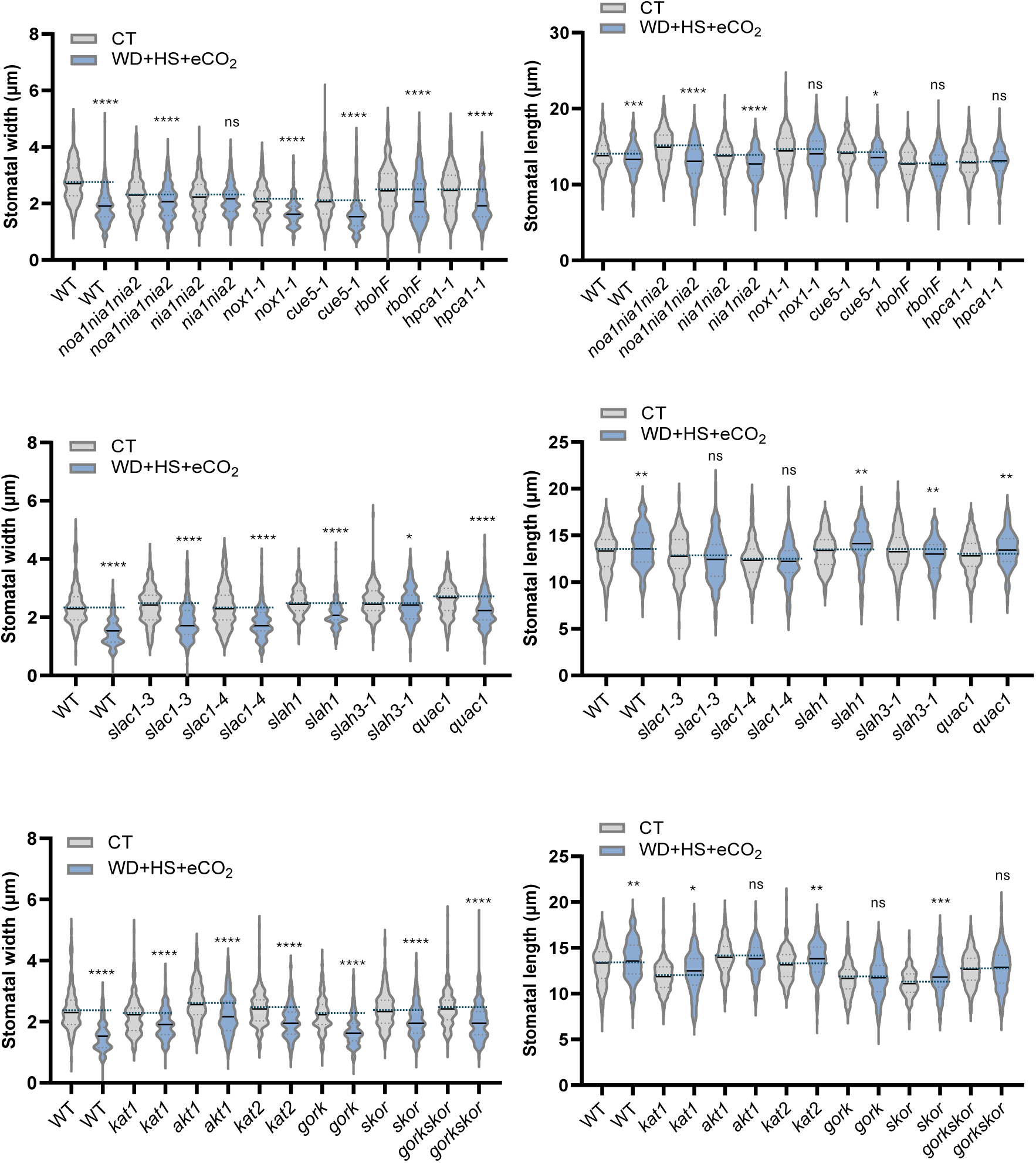
Measurements of stomatal width and length for the results shown in Figure 5. In support of Figure 5.

**Figure S8.**
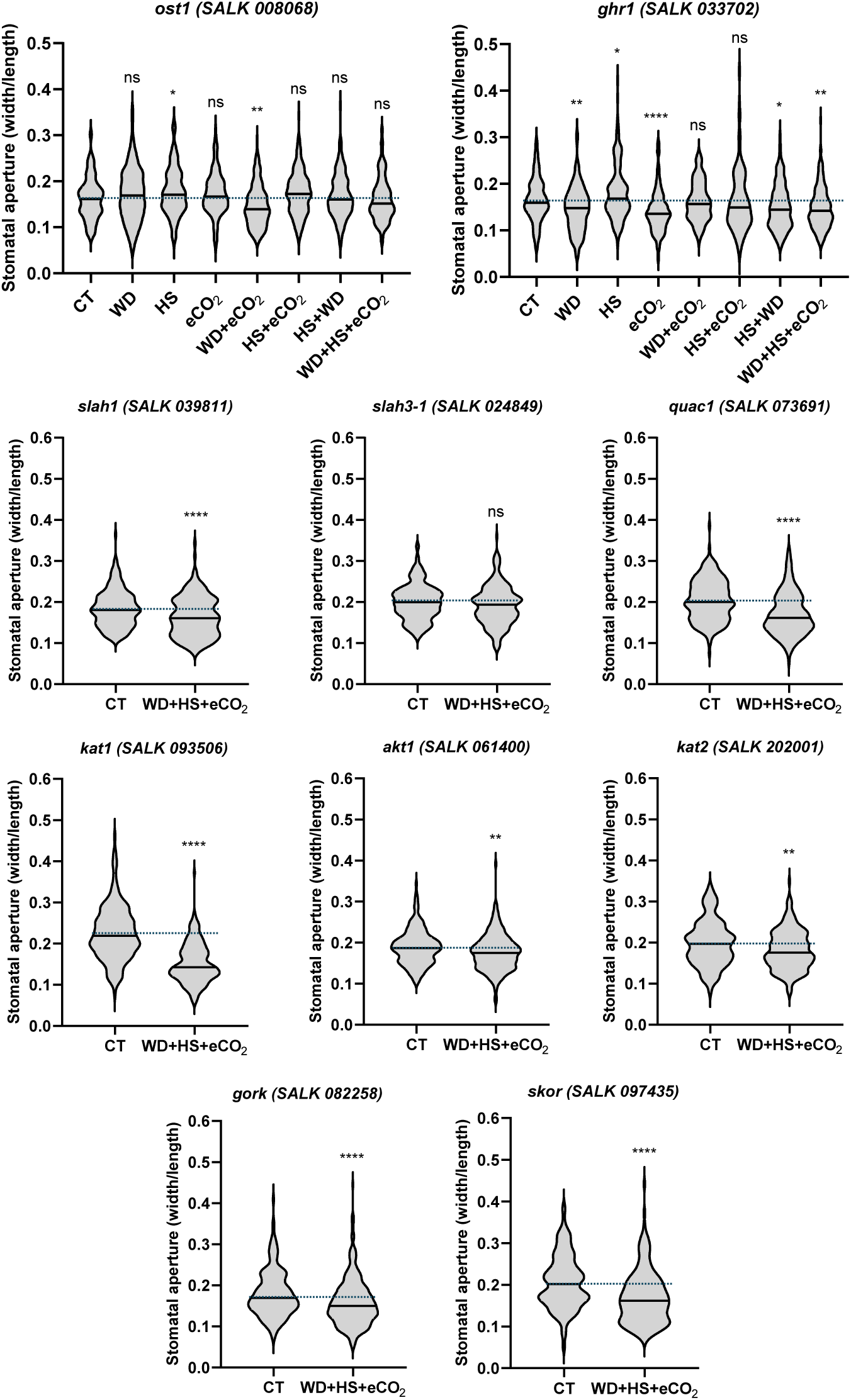
Stomatal aperture of different mutants (second alleles) grown under controlled growth conditions (CT), or subjected to water deficit (WD), heat stress (HS), and elevated CO_2_ (eCO_2_) in different combinations (In support of Figures 4, 5). One-way ANOVA followed by a Dunnett’s multiple comparisons test was used to determine significance. Asterisks denote statistical significance at **p < 0.01, ****p < 0.0001. Two-tailed Student’s t-test was used to determine significance at *p < 0.05. Details of the different mutants are given in the Supporting Information (Dataset S1). Abbreviations: K^+^ transporter 1 (akt1), elevated CO_2_ (eCO_2_), GUARD CELL HYDROGEN PEROXIDE-RESISTANT1 (ghr1), GUARD CELL OUTWARD RECTIFYING K+ CHANNEL (gork), heat stress (HS), CHANNEL IN ARABIDOPSIS THALIANA 1 (kat1), CHANNEL IN ARABIDOPSIS THALIANA 2 (kat2), non-significance (ns), OPEN STOMATA 1 (ost1), QUICK-ACTIVATING ANION CHANNEL 1 (quac1), stelar K+ outward rectifier (skor), SLOW ANION CHANNEL-ASSOCIATED 1 homologue 1 (slah1), SLOW ANION CHANNEL-ASSOCIATED 1 homologue 3 (slah3), water deficit (WD).

**Figure S9.**
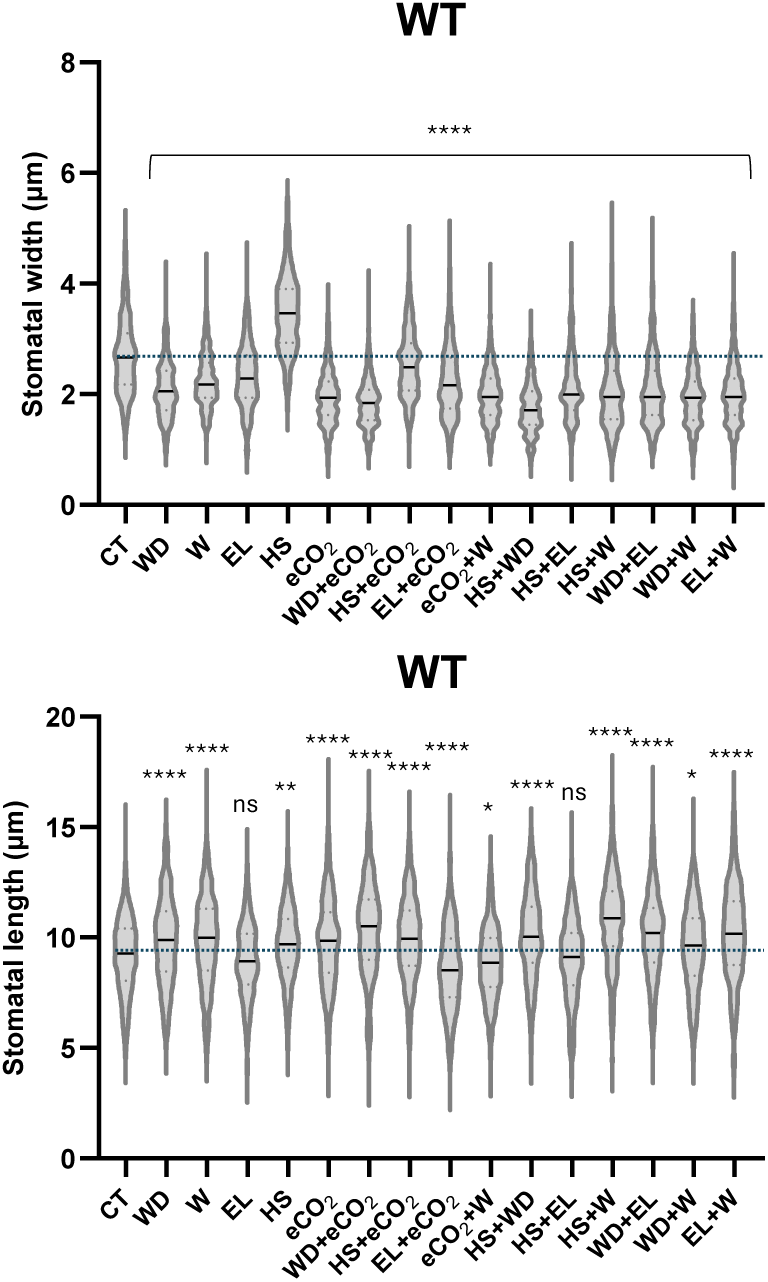
Measurements of stomatal width and length for the results shown in Figure 6a. In support of Figure 6.

## Notes

### Competing Interest Statement

The authors have declared no competing interest.

